# Common dandelion (*Taraxacum officinale*) leaf extract efficiently inhibits SARS-CoV-2 Omicron infection *in vitro*

**DOI:** 10.1101/2022.12.22.521558

**Authors:** Hoai Thi Thu Tran, Michael Gigl, Corinna Dawid, Evelyn Lamy

**Affiliations:** Molecular Preventive Medicine, University Medical Center and Faculty of Medicine, University of Freiburg, 79108 Freiburg, Germany; Food Chemistry and Molecular Sensory Science, Technical University of Munich, 85354 Freising, Germany

**Author notes:** Correspondence: Prof. Dr. Evelyn Lamy.

**Keywords:** COVID-19, *Taraxacum officinale*, SARS-CoV-2 prevention, Omicron variant, herbal medicine

## Abstract

As the COVID-19 pandemic continues to pose a health risk concern to humans, despite a significant increase in vaccination rates, an effective prevention and treatment of SARS-CoV-2 infection is being sought worldwide. Herbal medicines have been used for years and played a tremendous role in several epidemics of respiratory viral infections. Thus, they are considered as a promising platform to combat SARS-CoV-2. Previously, we reported that common dandelion (*Taraxacum officinale*) leaf extract and its high molecular weight compounds strongly suppressed *in vitro* lung cell infection by SARS-CoV-2 Spike D614 and Delta variant pseudotyped lentivirus. We now here demonstrate that *T. officinale* extract protects against the most prominent Omicron variant using hACE2-TMPRSS2 overexpressing A549 cells as *in vitro* model system. Notably, compared to the original D614, and the Delta variant, we could confirm a higher efficacy. Short-term interval treatment of only 30 min was then sufficient to block the infection by 80% at 10 mg/mL extract. Further subfractionation of the extract identified compounds larger than 50 kDa as effective ACE2-Spike binding inhibitors. In summary, the evolution of SARS-CoV-2 virus to the highly transmissible Omicron variant did not lead to resistance, but rather increased sensitivity to the preventive effect of the extract.

## Introduction

By November 2022, 634 million COVID-19 cases with 6.6 million deaths were confirmed worldwide [1], wherein, approximately 43 per cent of all infected patients experienced long term effect of infection (i.e. long COVID) [2]. Approximately 69% of the world’s population is vaccinated, meaning they have received the first dose, 60% are fully vaccinated (both doses), and a total of over 12 billion vaccine doses have been administered [3]. Although complete vaccination significantly reduces the likelihood of severe infection with SARS-CoV-2, some risk of viral transmission persists. Since the spike protein of the original D614 Wuhan strain has been used for the development of most currently available vaccines, the emergence and worldwide spreading of new SARS-CoV-2 variants with significant antigenic differences quickly resulted in a limited vaccine efficacy. The spike protein of the Omicron variant (B.1.1.529) has over 30 mutations with 15 mutations in the Spike receptor-binding domain (RBD) in comparison to the Wuhan strain. As a consequence, even fully vaccinated individuals are more frequently being diagnosed with COVID-19 as a result of breakthrough SARS-CoV-2 infection [4].

Herbal medicines have been used for centuries to treat epidemics and have generally been shown to be safe. Due to their distinct antiviral properties against various viruses, including SARS-CoV-2, they are being explored as interesting agents for complementary prevention and treatment of COVID-19 [5,6]. It has been suggested that the active compounds in these extracts could act through different modes of action, for instance, inhibition of protein-protein interaction as determined using computational screening assay, disrupting nuclear import and replication, or strengthening the immune system with reduced hyper-inflammatory responses [6,7]. However, the exact underlying mechanisms by which herbal medicines block viral infections remain to be investigated, as a number of active compounds with different molecular interactions are involved. To a certain extent, there have been some clinical trials using herbal medicines in combination with conventional therapies, which showed better results in terms of COVID-19 symptom relief and remission duration with higher lymphocyte counts [8,9]. Moreover, intranasal or intraoral administration of antiviral agents could be an important additional approach to prevent infection and spread of SARS-CoV-2 and other respiratory viruses at their entry sites, i.e., the nose and throat [10].

Recently, we demonstrated that the mechanism of action of *T. officinale* is to prevent the fusion of SARS-CoV-2 with human cells at the level of interaction of its spike proteins with the specific human cell surface enzyme angiotensin-converting enzyme 2 (ACE2). The potential use of monoclonal antibodies and the use of convalescent (antibody-rich) plasma in COVID-19 patients is also based on this mechanism [11]. Due to major concerns about immune evasion by the new viral variants and declining efficacy of available vaccines, we aimed to determine whether *T. officinale* extract could still provide efficacy against SARS-CoV-2 Spike Omicron variant infection. We also determined extract application times and further defined the active extract fraction.

## Results and Discussion

For the original SARS-CoV-2 virus utilization of the ACE2 receptor and other proteins has been confirmed as host cell entry, and this mechanism is still key for most emerging variants. Therefore, targeting Spike RBD – ACE2 binding is a promising approach to block virus entry. Our previous study showed that *T. officinale* extract could significantly suppress the binding between human ACE2 and SARS-CoV-2 Spike protein D614 as well as its four mutants [12]. Interestingly, the main bioactive chemicals responsible for the plant extract’s effects were in a high molecular weight fraction (≥ 5 kDa) based on our findings in both, a cell-free and cellular assay. Consequently, we further subfractionated the HMW fraction by using molecular weight cut-off filters of 5-10 kDa, 10-30 kDa, 30-50 kDa, and over 50 kDa. Figure 1 shows that in line with our previous study, the plant extract significantly reduced the Spike – ACE2 interaction already at 2 mg/mL by 24% and reached 54% at 50 mg/mL. Similarly, active compounds presented in the > 50 kDa HMW fraction showed a significant inhibition by 26% and 45% at the concentration equivalent to 2 and 50 mg/mL dried leaf extract, respectively. Whereas the other sub-fractions (5-10 kDa, 10-30 kDa, and 30-50 kDa) showed only minimal inhibition with less than 8%. This suggests that the effect is mainly caused by the > 50 kDa HMW fraction, but extensive research is still needed to identify the active compounds. We and others showed that the aqueous extract of *T. officinale* contains a variety of bioactive constituents, including more than 30 different phenolic compounds, sesquiterpene lactones, and polysaccharides [12,13]. For example, polyphenols have been attributed with a broad spectrum of antiviral activities, including anti-SARS-CoV-2 [14–16]. In a cell-free assay, polyphenols from hydroalcoholic pomegranate peel extract were shown to interfere with the spike-ACE2 interaction by binding to these proteins with a 10-fold higher affinity for the RBD domain. Among the polyphenolic compounds, punicalagin and ellagic acid were found to be the most active chemicals, and synergistic effects may exist [17]. However, using NMR analysis, we could confirm the absence of low molecular weight compounds in the HMW fraction. For sulfated polysaccharides (RPI-27 with a molecular weight of around 100 kDa derived from fucoidans), marked reduction of SARS-CoV-2 – ACE2 interaction at a non-cytotoxic concentration (IC50 of 8.3 ± 4.6 μg/mL) was demonstrated in Vero cells, and this may be due to multivalent binding to the S-protein of SARS-CoV-2 [15]. Using a docking model, the authors suggested that molecule binding to the S protein RBD was tighter with a larger oligosaccharide as compared to a lower molecular weight oligosaccharide

**Figure 1.**
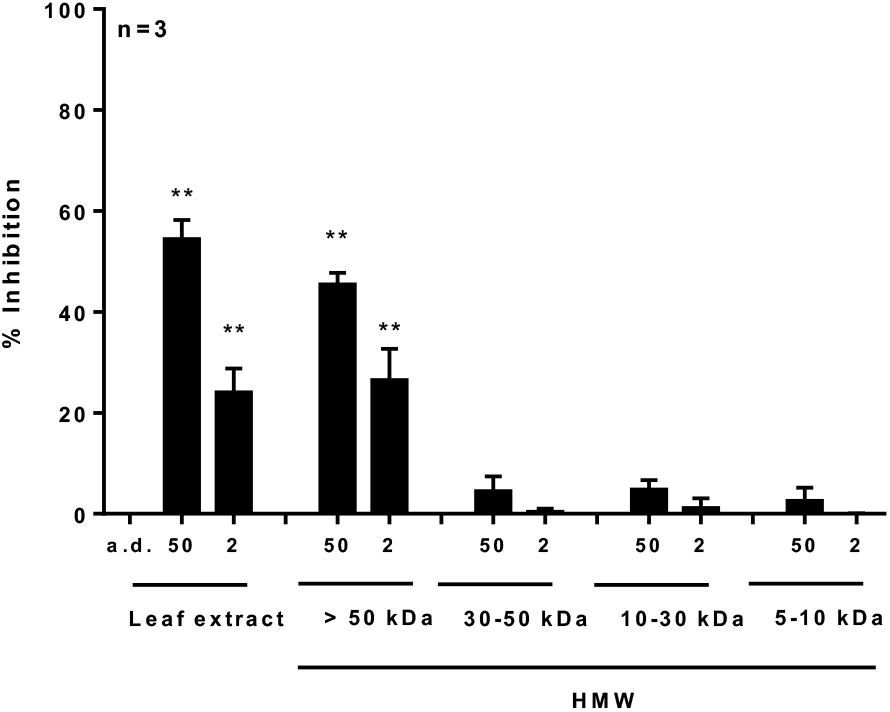
Effects of *T. officinale* high molecular weight subfractions on SARS-CoV-2 Spike ACE2 binding. Different concentrations of the plant extract or 5-10 kDa, 10-30 kDa, 30-50 kDa, and > 50 kDa HMW fractions (equivalent to dried leave weight) were used. The binding inhibition was determined using a SARS-CoV-2 Inhibitor Screening Kit. Bars are mean value + SD. Solvent control: distilled water (a.d.). Significance of difference was calculated relative to solvent control by one-way ANOVA followed by Bonferroni correction. * *p* < 0.05, ** *p* < 0.01.

*T. officinale* extract was shown previously by us to have a strong inhibitory effect on infection by SARS-CoV-2 Spike D614 and Delta (B.1.617.2) variant pseudotyped lentivirus [12]. Therefore, we here studied the efficacy of the extract to block B.1.1.529 Omicron variant transduction in A549-hACE2-TMPRSS2 cells using the same virus titer. Pre-treatment with the plant extract significantly inhibited viral transduction at 0.15 mg/mL by 24% and reached a peak (99%) inhibition at 2.5 mg/mL (Figure 2A). With this, the plant extract showed a much higher potency towards the SARS-CoV-2 Spike Omicron variant compared to the D614 (85 % inhibition at 20 mg/mL) and Delta variant (72% at 20 mg/mL), as reported previously [12]. Anti-hACE2 antibody (100 μg/mL) diminished viral infection by 82%, which was also slightly more effective compared to the results on the Delta variant (73%). Commonly, the standard test procedure normally provides test compound exposure for a few hours or even throughout the virus infection duration (up to 72 h) in cell-based models to identify a maximum possible treatment effect [17–19]. However, these treatment times are technically difficult or impossible to achieve in the clinical setting. A post-exposure prophylaxis that can be practiced on a daily basis would rather take place in several short treatment cycles. Based on these considerations, we repeated the experiment without *T. officinale* extract post-incubation. As a result, we still observed a substantial preventive effect against virus transduction, albeit with a lower inhibition efficacy (58% at 1.25 mg/mL, 99% at 5 mg/mL) (Figure 2B). This encouraged us to further test, whether only two rounds of short (30 min) extract treatment still could work. Interestingly, a preventive effect was already evident then at 2.5 mg/mL extract; 80% reduction in SARS-CoV-2 Spike Omicron particle infection was reached at 10 mg/mL (Figure 2C). As expected, extract efficacy (the half maximal effective concentration, EC50) of virus infection prevention increased with extension of extract treatment from EC50 of 5 mg/mL (30 min incubation before and after infection with washing in between), to 0.9 mg/mL (30 min pre-incubation prior 24 h transduction), to 0.5 mg/mL (30 min pre-treatment followed by 84 h incubation).

**Figure 2.**
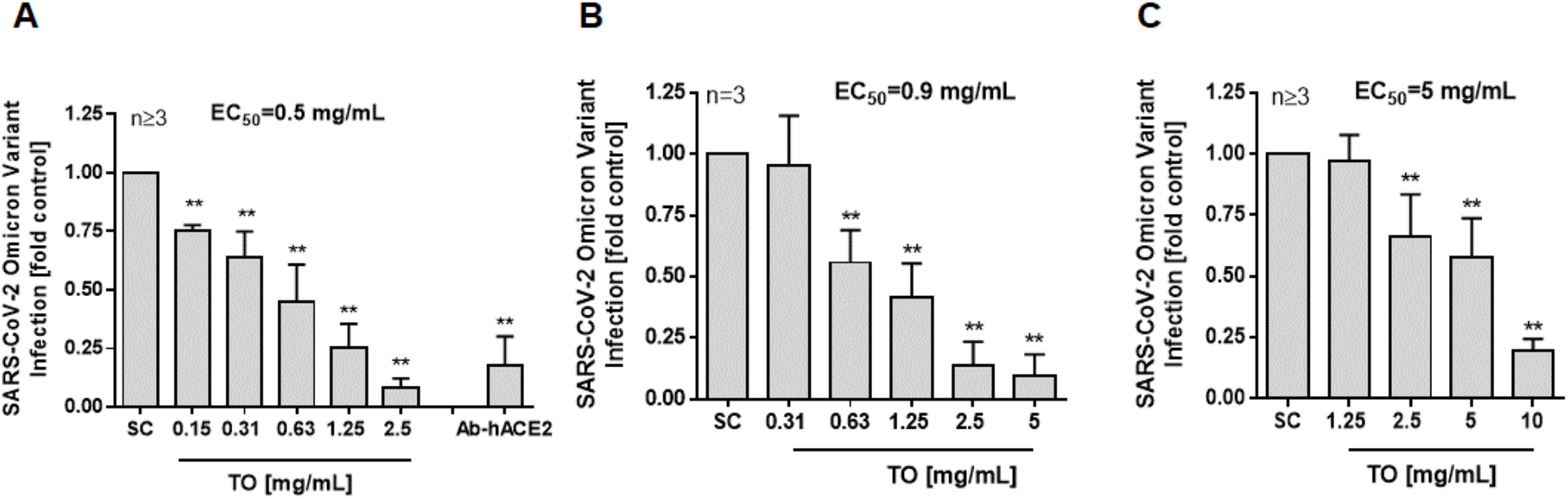
Inhibition of SARS-CoV-2 Spike B.1.1.529 Omicron variant by *T. officinale* extract. A and B) Cells were pre-treated with *T. officinale* (TO) extract for 30 min before transfection with 7500 TU/mL SARS-CoV-2 Spike Omicron variant virus particles for 24 h. The cells were post-incubated in fresh medium (A) containing TO for 60 h or (B) without TO for 48 h. C) Cells were exposed to TO for 30 min, then washed with PBS, and fresh medium containing 7500 TU/mL SARS-CoV-2 Spike Omicron variant virus particles was added for 24h. Cells were treated again for 30 min with TO in fresh medium, washed 1× with PBS, and post-incubated for another 48 h. Luminescence was measured after 15 min. Solvent control (SC): 10% distilled water. Inhibitor positive control: 100 μg/ml anti-hACE2 antibody. Data are mean value + SD. Significance of difference was calculated relative to respective solvent control by oneway ANOVA followed by Bonferroni correction. * *p* < 0.05, ** *p* < 0.01.

According to the German Commission E and the European Scientific Cooperative for Phytotherapy (ESCOP) [20], the recommended daily intake of the plant is up to 8 g (i. e. 20 – 30 mg/mL). This makes it interesting for an intervention in the oral cavity, as it would be realistic and safe to conduct short, 30 min interval treatments in expectation or after potential Sars-CoV-2 virus exposure. In our previous experiments we already demonstrated at least 30 min stability of the bioactive compounds in human saliva at body temperature [12]. The oral cavity, as an essential part of the upper digestive tract, plays a key role in the transmission and pathogenicity of SARS-CoV-2. Moreover, the new Omicron virus variants appear to infect the gastrointestinal tract in addition to the respiratory tract. Gastrointestinal symptoms sometimes occur before respiratory symptoms [21,22], and compared to the lung, even higher expression levels of the ACE2 receptor were found in the gastrointestinal tract [23].

## Materials and Methods

### Plant material and extraction

*T. officinale* dried leaves were purchased from Achterhof (Uplengen, Germany; batch number B370244). Plant extraction was done as described before [12]. Briefly, 100 mg/mL plant extract in distilled water was incubated for 1 h at room temperature (RT). The aqueous extract was collected and centrifuged at 16000 × g, 3 min, RT. Subsequently, the supernatant was filtered through a 0.22 μm filter before use.

### Isolation and purification of high molecular weight (HMW) fractions from plant material via sequential ultrafiltration

First, the plant extract was prepared by submerging dried plant material (200 g) in water (2 L) followed by incubation in the dark at RT for 1 h. After filtration with a suction filter, the resulting extract was subjected to a sequential ultrafiltration procedure using a crossflow system (Sartorius Stedim Biotech, Göttingen, Germany). Filtration was carried out with four molecular weight cut-off filters (50 kDa, 30 kDa, 10 kDa, and 5 kDa) resulting in the following fractions: >50 kDa, 30–50 kDa, 10–30 kDa, and 5–10 kDa. Purification of HMW fractions was achieved by flushing with water (5 L each) to remove low molecular weight compounds from the extract. Purification was verified by nuclear magnetic resonance (NMR) spectroscopy as reported previously [24] The yields of all isolated fractions were determined by weight after lyophilization and stored at −20°C until use.

### SARS-CoV-2 Spike–ACE2 Interaction Inhibition

To determine the impact of the plant extract and its fractions on SARS-CoV-2 Spike ACE2-binding, SARS-CoV-2 Inhibitor Screening Kit (Fisher Scientific GmbH, Schwerte, Germany) was used according to the manufacturer’s instructions and as described previously [12].

### A549-hACE2-TMPRSS cell culture

Human lung A549 cells, stably expressing hACE2 and TMPRSS2 (A549-hACE2-TMPRSS2), were obtained from InvivoGen SAS (Toulouse Cedex, France). The cell culture was done according to the manufacturer’s protocol in a DMEM medium containing 4.5 g/L glucose, 2 mM L-glutamine, 10% heat-inactivated fetal bovine serum (FBS), 100 U/mL penicillin/streptomycin, 100 μg/mL normocin, 0.5 μg/mL puromycin, and 300 μg/mL hygromycin. The cells were cultured in a humidified incubator at 37°C, 5° CO2 air atmosphere at 37°C.

### SARS-CoV-2 Spike B.1.1.529 (Omicron) Variant transduction assay

SARS-CoV-2 Spike B.1.1.529 (Omicron) variant pseudotyped lentivirus particles were purchased from Biomol (Hamburg, Germany). The transduction assay was performed as described previously [12]. Briefly, 1 × 10^5^ cells/cm^2^ (< 15 passages) were seeded in 96-well plates in DMEM medium, 4.5 g/L glucose, supplemented with 2 mM L-glutamine, 10% heat-inactivated FBS. After 8 h, the cells were either a) pre-treated with plant extract or anti-hACE2 antibody for 30 min before 24 h lentiviral transduction, followed by 60 h post-incubation in fresh medium containing plant extract; b) infected with lentivirus particles for 24 h before 48 h post-incubation without plant extract; or c) treated with plant extract for 30 min, and then washed with warm PBS, before infection with lentivirus particles for 24 h. After that, the cells were exposed to a fresh medium containing plant extract for another 30 min, washed with warm PBS again, and finally, cells were incubated for another 48 h in fresh medium. Luminescence (one-step luciferase assay system, Biomol) was detected after 15 min using a multi-plate reader (Tecan Group Ltd, Crailsheim, Germany). 10% filtered distilled water was used as solvent control (SC).

### Statistical Analysis

Data were analyzed using GraphPad Prism 6.0 software (La Jolla, CA, USA). Results are depicted as means + SD. Significance of difference was calculated relative to solvent control (10% distilled water) by the one-way ANOVA test followed by Bonferroni correction. P values < 0.05 (*) were considered statistically significant and < 0.01 (**) were considered highly statistically significant.

## Conclusions

The findings confirm our previous observations and provide supportive evidence on the potential benefits of *T. officinale* extract and its > 50 kDa HMW fraction against SARS-CoV-2 wildtype as well as variants of concern via interfering with Spike RBD – ACE2 binding. Still, the mechanism for infection prevention, and also the active compounds responsible for it, are not yet fully understood and require further evaluation. Given the rapid emergence of new mutations and the prevalence of the Omicron variant, which has been shown to decrease the prophylactic and therapeutic efficacy of neutralizing antibodies by more than a hundredfold [25,26], there is an urgent need for clinical trials of agents that exhibit potent anti-SARS-CoV-2 Omicron variant infection. Therefore, *T. officinale* is worth further clinical research to determine its supportive prevention and therapy.

## Author Contributions

Conceptualization, E.L. and H.T.T.T; methodology, H.T.T.T and M.G.; validation, E.L., H.T.T.T, M.G., and C.D.; formal analysis, H.T.T.T; investigation, H.T.T.T and M.G; resources, E.L. and C.D.; writing—original draft preparation, H.T.T.T and E.L.; writing— review and editing, E.L., H.T.T.T, M.G., and C.D.; supervision, E.L. and C.D.; funding acquisition, E.L. and C.D. All authors have read and agreed to the published version of the manuscript.

